# Quantifying Brittle Crack Opening in Human Trabecular Bone Using Synchrotron XCT–DVC

**DOI:** 10.64898/2026.03.24.714043

**Authors:** Dhruv Vasooja, Ahmet Cinar, Mahmoud Mostafavi, James Marrow, Christina Reinhard, Ulrich Hansen, Richard Leslie Abel

**Affiliations:** MSK Laboratory, Department of Surgery and Cancer, Faculty of Medicine, Imperial College London, W12 0BZ UK; Department of Materials, University of Oxford, Parks Road, Oxford, OX13PH UK; Frontier Robotics, The National Robotarium, Heriot-Watt University, EH14 4AS UK; Department of Mechanical and Aerospace Engineering, Monash University, 14 Alliance Lane, Clayton 3800, Australia; The University of Manchester, Faculty of Science and Engineering, Oxford Road, Manchester, M13 9PL UK; The University of Manchester at Harwell, Diamond Light Source, Harwell Science & Innovation Campus, Chilton, OX11 0DE UK; Department of Mechanical Engineering, Faculty of Engineering, Imperial College London, London, SW7 2AZ UK

## Abstract

**Introduction:** Trabecular bone exhibits brittle behaviour governed by microscale deformation and damage processes, yet quantitative characterisation of crack progression remains challenging because classical fracture mechanics approaches do not apply to architecturally discontinuous porous tissues. This study evaluates whether synchrotron X-ray computed tomography (XCT) combined with digital volume correlation (DVC) can provide a practical experimental approach for quantifying crack opening behaviour in human trabecular bone.

**Method:** Semicylindrical specimens harvested from femoral heads of hip-fracture donors (n = 5) and non-fracture controls (n = 5) underwent stepwise three-point-bending during XCT imaging. Full-field displacement maps enabled direct measurement of crack mouth opening displacement (CMOD), crack length (a), and their ratio, CMOD/a, used here as a geometry-normalised comparative descriptor of brittle response. Automated crack segmentation using phase-congruency crack detection (PCCD) was compared against manual measurements.

**Results:** XCT-DVC successfully resolved three-dimensional displacement discontinuities during crack initiation and propagation in all specimens. Hip-fracture donors exhibited significantly lower critical crack-opening ratios (CMOD/*a*)* than Controls (0.31 vs 0.47; p = 0.008) and reached mechanical instability at lower applied loads, consistent with a more brittle structural response under this test configuration. Despite these differences, total crack extension (Δ*a*\*) was similar between groups. Automated crack tracking using phase-congruency–based segmentation showed excellent agreement with manual measurements (*r²* = 0.98), confirming reliable extraction of crack geometry from DVC displacement fields.

**Discussion:** These results indicate that XCT-DVC can provide a practical approach for quantifying crack-opening behaviour in trabecular bone when classical fracture-mechanics parameters are not applicable in anatomically constrained specimens. The reduced critical crack-opening ratios and earlier instability observed in Hip-fracture donors are consistent with a more brittle comparative mechanical response that is not captured by crack extension alone. The strong agreement between automated and manual crack measurements further supports displacement-based descriptors as reliable comparative indicators of brittle behaviour in porous, architecturally discontinuous tissues.

Graphical abstract

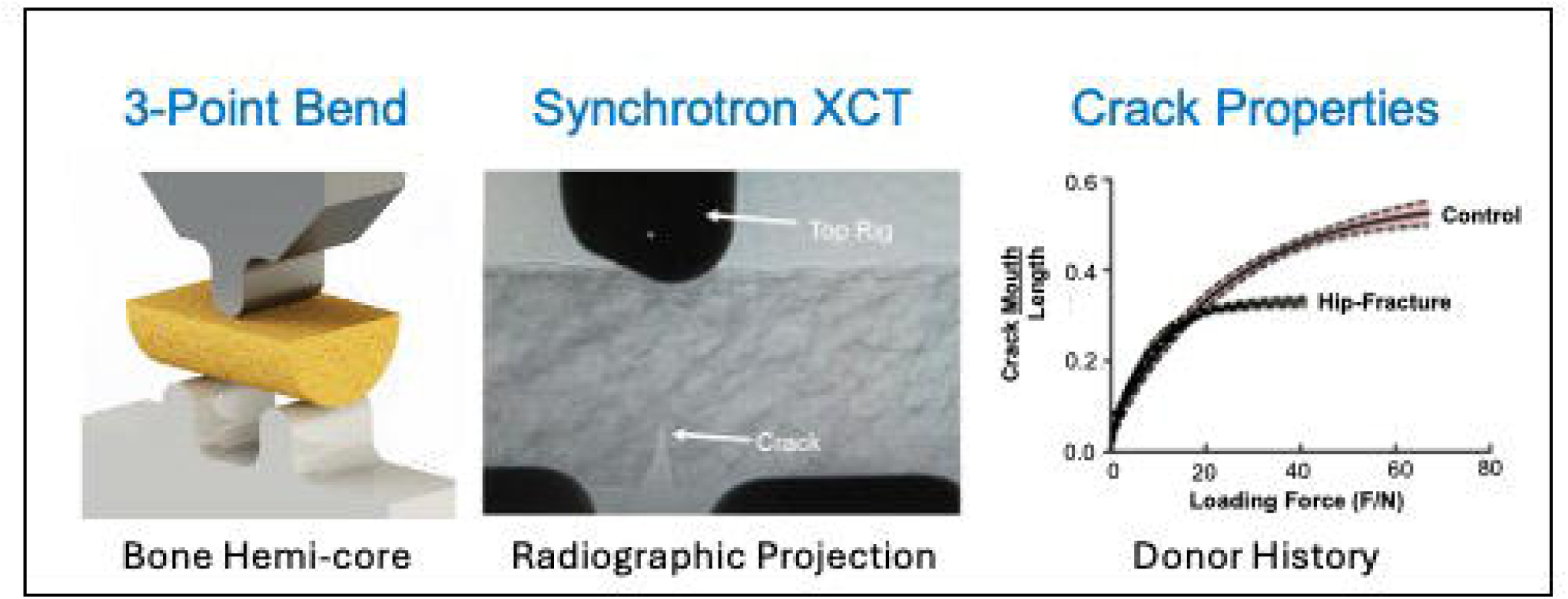

## 1. Introduction

Trabecular bone is a hierarchical, porous solid that fails by the nucleation, growth, and coalescence of cracks within a discrete lattice of struts and plates. From a fracture-mechanics point of view, it behaves as a quasi-brittle cellular material rather than a homogeneous continuum, with failure governed by local microstructural interactions at the crack tip and along the crack path. Classical fracture-toughness approaches, which assume a macroscopically continuous medium and a single, well-defined crack front, therefore cannot provide reliable stress-intensity (K) or J-integral values for small trabecular volumes from anatomical sites such as the femoral head, necessitating alternative, displacement-based measures of crack opening. Consequently, there has been a broad transition from global metrics such as ultimate strength toward localised, microstructure-resolved measures of deformation, crack opening and damage accumulation, enabled by micro-CT, image correlation and related high-resolution techniques (Cook & Zioupos, 2009; Gillard et al., 2014; Palanca et al., 2016; Turunen et al., 2020; Yan et al., 2020; Rowe et al., 2025). Despite these advances, it is still unknown whether full-field displacement measurements can provide consistent, geometry-normalised descriptions of crack opening during fracture in trabecular bone—an architecturally discontinuous tissue for which classical toughness approaches do not apply.

A wide range of experimental approaches has been used to probe the fracture behaviour of cancellous and cortical bone (Ritchie et al., 2008; Cook & Zioupos, 2009), including micro-scale fracture tests (Zimmermann et al., 2019) and macro-scale fracture tests on notched specimens (Ritchie et al., 2008), high-resolution imaging of microdamage (Jin et al., 2017; Ma et al., 2017), and computational models based on CT-derived micro-structures (Niebur et al., 2000; Keaveny et al., 2001; Morgan et al., 2003). Local fracture properties have been estimated using J–R curves, energy-based measures and crack-tip opening parameters (Cook and Zioupos, 2009; Zimmermann et al., 2019) often in combination with digital image correlation (DIC) at the specimen surface (Yan et al., 2020). However, in trabecular bone the strongly discontinuous architecture (Keaveny et al., 2001), the limited specimen size available for testing (Cook & Zioupos, 2009), and the tendency for cracks to twist and branch through the 3D lattice (Barak et al., 2018; Turunen et al., 2020) make it difficult to define a unique crack front or to apply standard linear elastic fracture mechanics (LEFM) assumptions (Cook & Zioupos, 2009; Keaveny et al., 2001).

Digital volume correlation (DVC) with Synchrotron X-ray computed tomography (XCT) offers a route around some of these limitations. The high resolution of XCT facilitates full-field 3D displacement and strain measurements inside trabecular bone specimens. Early DVC studies in bone established accuracy and precision for trabecular architectures, showing that appropriately chosen sub-volume sizes can resolve internal strain fields and validate finite element models of vertebral and cancellous specimens (Liu and Morgan, 2007; Gillard et al., 2014; Palanca et al., 2016; Dall’Ara et al., 2017). More recent work has used DVC to examine sub-trabecular strain evolution, demonstrating that regions about to crack experience markedly higher local strains than the bulk, and that tissue-level strain thresholds for damage are higher than previously assumed (Turunen et al., 2020; Yan et al., 2020). These studies highlight the importance of local kinematics and damage processes in governing apparent brittle behaviour, but they do not directly provide a geometry-controlled description of crack opening during unstable fracture.

In parallel, a substantial XCT–DVC literature has developed in engineering quasi-brittle materials, where many of the same mechanical challenges arise. In nuclear graphite and other porous solids, XCT combined with DVC has been used to map full-field displacements and crack-tip strain fields during stable crack growth, revealing non-linear process zones and cohesive fracture behaviour (Mostafavi et al., 2013; Barhli et al., 2017; Wade-Zhu et al., 2020). The DVC-measured displacement field has been exploited to reconstruct three-dimensional crack opening displacement (COD) profiles, to estimate effective crack lengths, and to calibrate cohesive-zone models, providing displacement-based comparative indicators of brittle behaviour even when conventional *K_IC_* measurements are problematic (Mostafavi et al., 2013; Barhli et al., 2017). Similar strategies are now being extended to other quasi-brittle systems such as concrete and hydrogels, underlining the generality of displacement-based fracture descriptors derived directly from 3D imaging (Oesch et al., 2020).

A key technical development has been the use of phase-congruency-based crack detection to identify displacement discontinuities directly from DVC fields. By combining DVC with phase congruency analysis, crack surfaces can be segmented objectively in three dimensions, allowing crack shape, size and opening to be quantified with minimal operator bias. This methodology has been applied in graphite and other heterogeneous materials to extract local fracture parameters from 3D data (Cinar et al., 2017; Barhli et al., 2017). More recently, similar ideas have been adapted to bone. Trabecular bone fracture tests have used XCT–DVC with phase congruency to identify 3D cracks and, via finite element back-analysis, to estimate critical stress intensity factors without strong dependence on specimen geometry or direct load measurements (Yan et al., 2020). In cortical bone, automatic crack detection based on displacement discontinuities has been used to generate crack mouth opening displacement (CMOD)–load fracture resistance curves and to relate crack-tip strains and tortuosity to microstructural features (Cinar et al., 2017).

Despite this progress, there is still no commonly adopted laboratory approach for characterising brittle crack opening in human trabecular bone within the geometric limitations imposed by small, anatomically derived specimens. In cancellous bone, most studies have focused on (i) bulk strength and stiffness, (ii) local strain patterns in intact trabeculae, or (iii) specimen-specific fracture-toughness estimates obtained through complex inverse analyses (Cook & Zioupos, 2009; Yan et al., 2020; Turunen et al., 2020). However, these approaches either do not track crack evolution directly or rely on specimen geometries incompatible with trabecular architecture. What is lacking is a comparatively simple, geometry-normalised descriptor of crack opening—one that reduces sensitivity to specimen size and crack-front geometry and can be extracted directly from XCT-DVC data to enable meaningful comparison of brittle behaviour across donors, material conditions, and disease states.

The overarching aim of this study was to test whether synchrotron XCT combined with DVC can provide a geometry-normalised, displacement-based approach for quantifying brittle crack opening in human trabecular bone. We addressed this aim through two specific objectives. First, we used XCT–DVC during three-point bending of trabecular semi-cores to quantify crack mouth opening displacement (CMOD), crack length a, and the geometry-normalised crack-opening ratio CMOD/a throughout crack propagation. Second, we tested whether these CMOD-based measures, and their critical values at the onset of instability, can act as robust specimen-scale descriptors of brittle behaviour that distinguish trabecular bone from hip-fracture donors and from non-fracture controls. We also evaluated whether these quantities can be extracted reliably using automated phase-congruency–based crack segmentation compared against manual measurements. We hypothesised that trabecular bone from hip-fracture donors would exhibit lower critical (CMOD/a)* values at the onset of instability than bone from non-fracture controls, consistent with a more brittle comparative mechanical response.

## 2. Materials and Methods

This study combined stepwise in situ three-point bending with synchrotron XCT to quantify brittle crack opening in trabecular bone. The analysis pipeline comprised: (i) reconstruction and preprocessing of tomographic volumes; (ii) computation of full-field 3D displacement fields using digital volume correlation (DVC); (iii) quantification of crack geometry using both manual inspection of resliced XCT image stacks and automated phase-congruency–based crack detection (PCCD) applied to DVC displacement discontinuities; and (iv) derivation of displacement-based crack-opening descriptors. The primary descriptors were crack mouth opening displacement (CMOD), defined as the normal displacement separation across the notch mouth from DVC; crack length (a), measured from the notch root along the detected crack path; and the geometry-normalised crack-opening ratio CMOD/a. The critical plateau value (CMOD/a)* was taken at the onset of unstable crack growth, and total crack extension was quantified as Δa*. Owing to the deviation from standard fracture-mechanics configurations and the short span relative to specimen depth, the loading condition involved mixed-mode components; classical fracture-toughness parameters (e.g. KIC, J) were therefore not evaluated, and interpretation is restricted to comparative, specimen-scale CMOD-based descriptors of brittle response. CMOD/a is used here as a geometry-normalised comparative descriptor of crack-opening behaviour, not as an intrinsic material property or a substitute for formal fracture-toughness parameters.

### 2.1 Specimen procurement and preparation

A total of 10 trabecular bone specimens were obtained from femoral heads of two donor groups: hip-fracture patients (Hip-Fx) and healthy ageing non-fracture controls (Control) (n = 5 each). Hip-Fx femoral heads were collected from patients undergoing arthroplasty for femoral neck fracture at Imperial College Healthcare NHS Trust, London, UK, while non-fracture control femoral heads were obtained from cadaveric donors at the Vesalius Clinical Training Centre, University of Bristol (as in Jin et al., 2017; Ma, 2018). Donor characteristics are summarised in Table 1. All procedures complied with the Declaration of Helsinki; the Imperial College Tissue Bank granted ethical approval (Reference R13004), and all patients/donors or their representatives provided informed consent for research use of tissue. Individuals with primary bone disease, metastatic malignancy, or other causes of secondary bone disease were excluded.

**Table 1.**
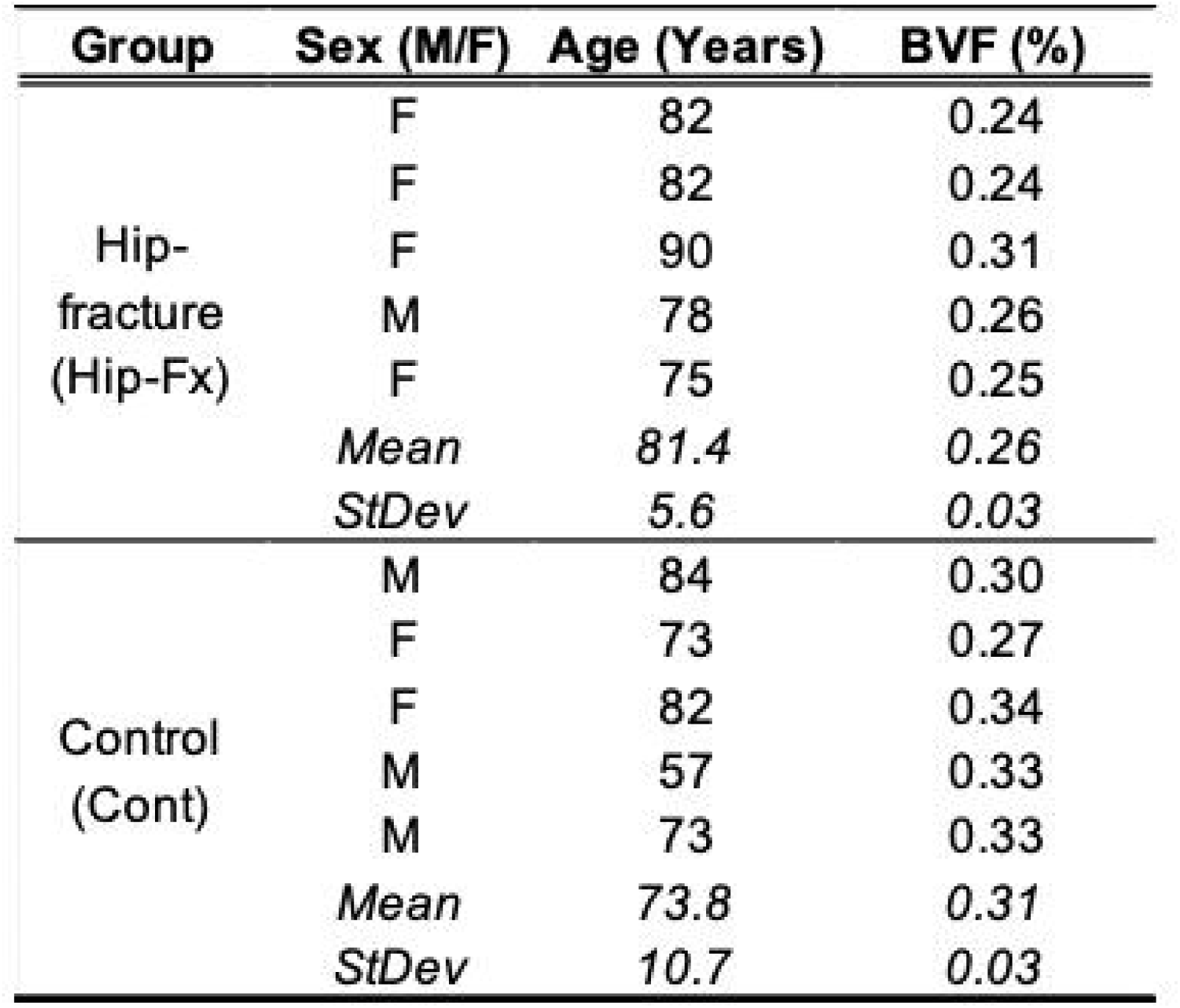
Donor and trabecular bone characteristics for hip-fracture and control groups. Demographic and structural parameters for trabecular bone specimens obtained from the femoral heads of donors with fragility hip-fracture (Hip-Fx) and age-comparable non-fracture controls (Cont). Hip-Fx donors tended to be older (∼7.6 years, p = 0.208) and exhibited significantly lower bone volume fraction (BV/TV; −20%, p = 0.019). BV/TV represents the proportion of total volume occupied by mineralised bone tissue. M = male; F = female.

| **Group** | **Sex (M/F)** | **Age (Years)** | **BVF (%)** |
| --- | --- | --- | --- |
| Hip-fracture<br>(Hip-Fx) | F | 82 | 0.24 |
|  | F | 82 | 0.24 |
|  | F | 90 | 0.31 |
|  | M | 78 | 0.26 |
|  | F | 75 | 0.25 |
|  | *Mean* | *81.4* | *0.26* |
|  | *StDev* | *5.6* | *0.03* |
| Control<br>(Cont) | M | 84 | 0.30 |
|  | F | 73 | 0.27 |
|  | F | 82 | 0.34 |
|  | M | 57 | 0.33 |
|  | M | 73 | 0.33 |
|  | *Mean* | *73.8* | *0.31* |
|  | *StDev* | *10.7* | *0.03* |

From each femoral head, cylindrical trabecular cores were prepared from the primary compressive trabecular arcade in the region directly superior to the trabecular chiasma, following our previous micro-CT and mechanical studies (Jin et al., 2017; Ma, 2018). Cores (10 mm height × 7 mm diameter) were drilled using a low-speed (500 rpm) bench pillar drill (Jet JDP-15B, Jet Tools, La Vergne, TN, USA) under constant irrigation with saline using a diamond coring bit (DK Holdings Ltd, Staplehurst, UK) to minimise thermal and mechanical damage. The coring trajectory was aligned with the principal load-bearing trabecular direction to ensure anatomical and mechanical comparability across donors.

Cores were sectioned to form a semi-cylindrical test specimen (semi-core) following Ma (2018) (Figure 1). Each cylindrical core was halved longitudinally using a low-speed diamond wafering saw (Isomet, Buehler, Germany) to generate a semicylindrical specimen. The final semi-core geometry was 7 mm (thickness at the top surface) × 3.5 mm (radius/height) × 12 mm (length), representing the largest specimen that could be obtained from the available material while approximating ASTM E1820 constraints (Ma, 2018). A sharp, mode-I starter notch was introduced at the mid-span of the top (putative tensile) surface using a razor blade, producing a notch length of approximately 0.8 mm and width of 0.15 mm. The notch was oriented perpendicular to the loading direction and centred along the longitudinal axis of the semi-core.

**Figure 1.**
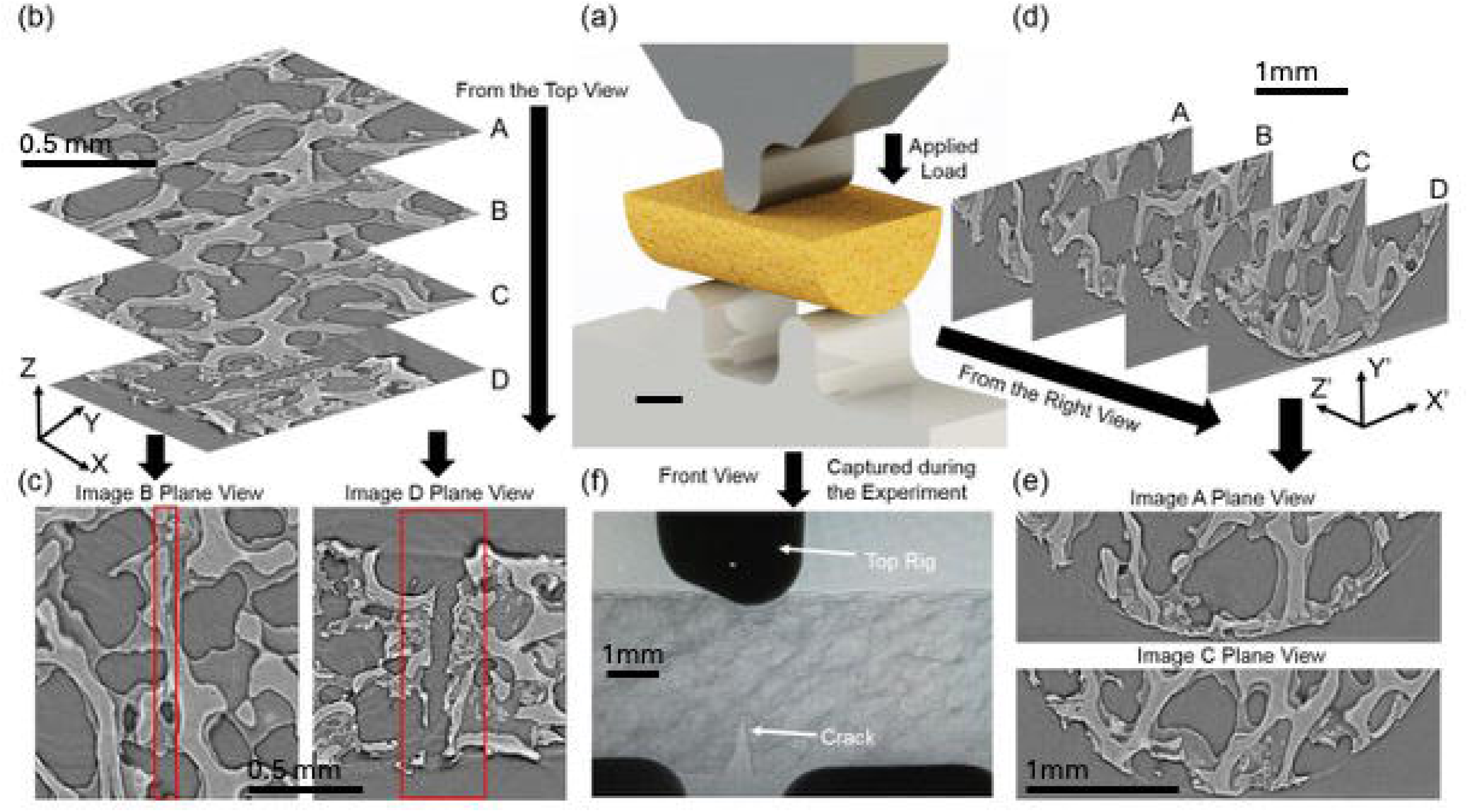
Synchrotron XCT imaging of trabecular bone during in situ three-point-bending. (a) Three-point-bending experimental setup used during synchrotron imaging at beamline I12. (b) Volume renderings (top view) showing trabecular microarchitecture. (c) Top view images highlighting the crack tip (Image B) and the crack mouth (Image D). (d) Resliced volume images from the right view perspective. (e) High-resolution detail of trabecular architecture from the right view. (f) Radiographic projection showing the notched crack during loading. Images acquired at 3.2 μm voxel size.

**Figure 2.**
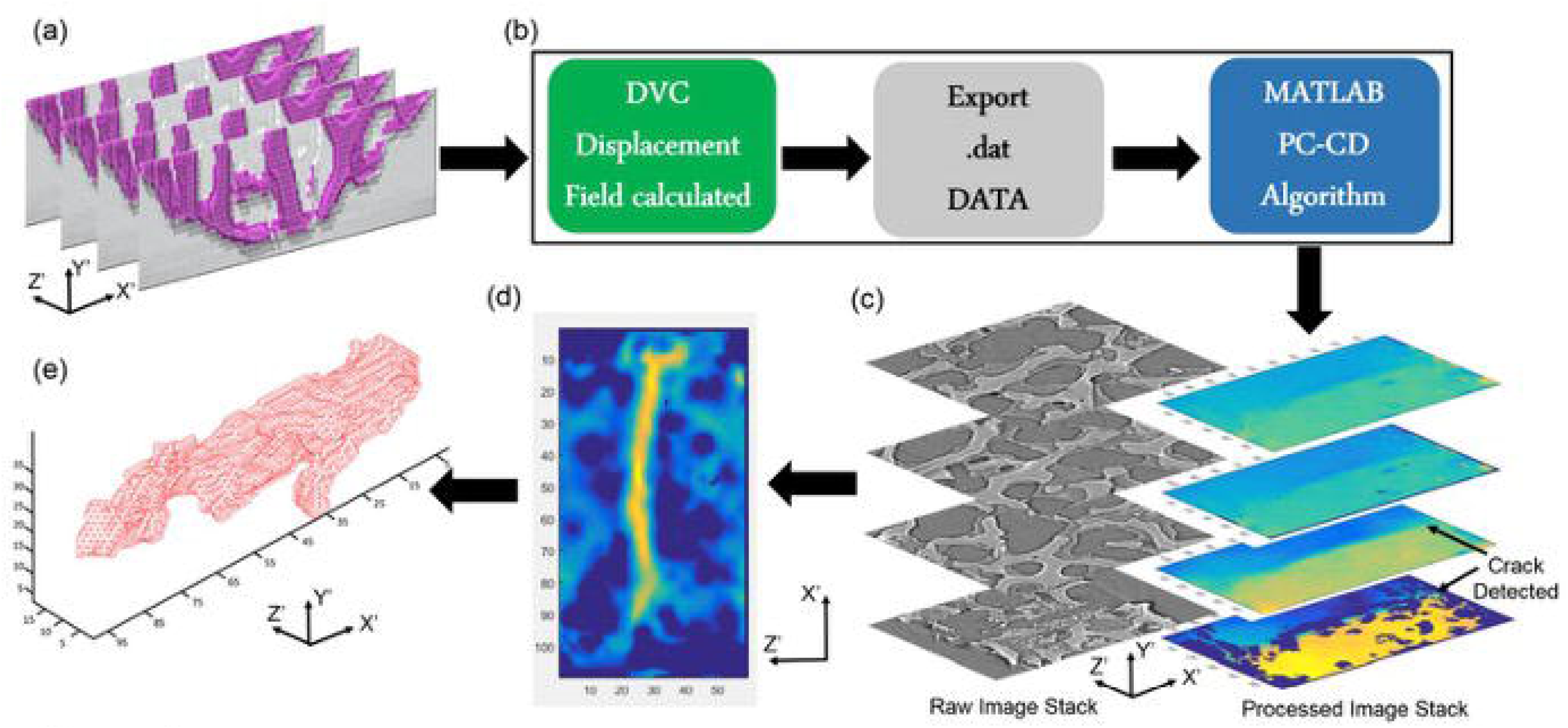
Automated crack detection using the Phase Congruency Crack Detection (PCCD) algorithm. (a) DVC-derived displacement field (right view) indicating localised discontinuities. (b) Processing workflow from displacement fields through PCCD segmentation. (c) 3D crack detection in the Y′ direction, aligned with trabecular architecture. (d) Discontinuity field visualisation in the X′Z′ plane showing selected seeding points. (e) Final 3D segmented crack surface used for quantitative crack length analysis with sub-voxel precision.

**Figure 3.**
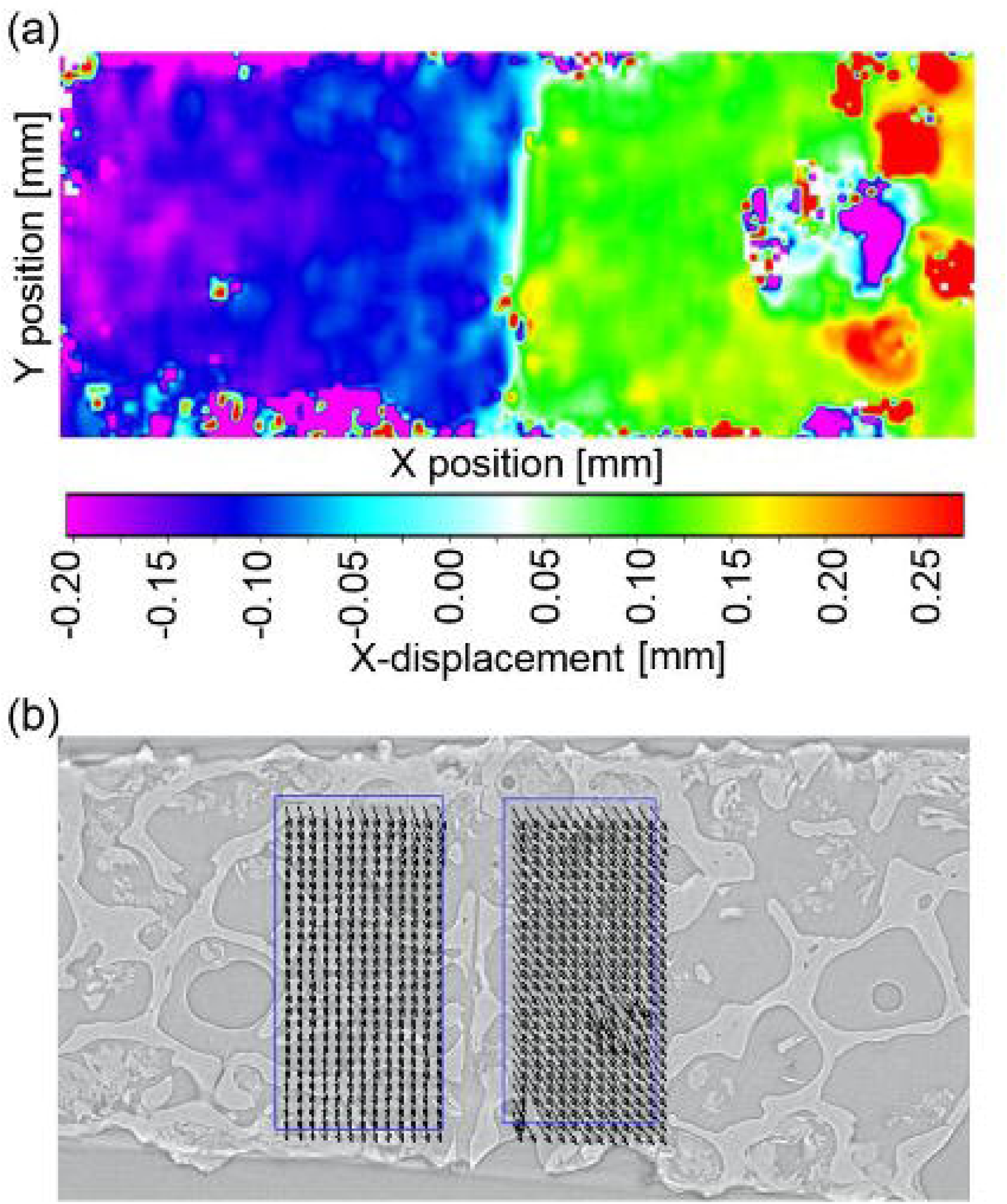
Quantification of crack mouth opening displacement (CMOD) from DVC. (a) X-direction displacement contour plot showing a clear displacement discontinuity across the crack interface (units: mm). (b) Application of geometric masks on opposing crack faces to extract displacement separation for CMOD calculation; vectors represent local displacement direction and magnitude.

In total, 10 semi-core specimens were prepared for fracture testing (n = 5 Hip-Fx; n = 5 Control), corresponding to one semi-core per femoral head. After preparation, cores and semi-cores were wrapped in PBS-soaked gauze and stored at −80 °C and only removed immediately prior to imaging and testing. During thawing, handling, and in situ testing, specimens were kept hydrated in PBS buffer solution. Prior to imaging, specimens were sprayed with PBS buffer solution and placed in a thin-walled plastic tube to limit specimen dehydration during repeated scans. Experiments were conducted at room temperature.

### 2.2 Mechanical testing and synchrotron XCT imaging

Mechanical loading and imaging were performed in situ at the Joint Engineering, Environmental and Processing (JEEP) beamline I12, Diamond Light Source (Harwell, UK), during beamtime allocation EE11204-1, using a custom three-point bending rig designed for small, irregular bone specimens (Ma, 2018; Yan et al. 2020). Semi-core specimens were mounted with the curved surface resting on two bottom support rollers (radius 2.0 mm) and the machined, notched flat surface facing upwards under the central loading roller. The support span between the two bottom rollers was 8 mm, selected to maximise bending stresses in the region of interest while accommodating the limited specimen length. Prior to imaging, each specimen was carefully aligned within the rig so that the notch lay at mid-span and within the synchrotron field of view.

Loading was applied in displacement control at a fixed nominal strain rate of 0.001 s−1 using a 1 kN load cell (1 N resolution), following Ma (2018). A preload of 2 N was applied to stabilise specimen positioning and to acquire a reference (unloaded) tomographic volume without specimen movement. Load was then increased in 5 N increments up to failure. At each increment, the actuator was paused, load held constant, and a full XCT scan acquired before proceeding to the next increment, following the protocol of Ma et al. (2018) and Yan et al. (2020). A total of 12 loading stages (including the reference state) were targeted where possible; in specimens that failed earlier, imaging continued until unstable crack propagation or loss of structural integrity precluded further loading (typically 9–12 stages). The applied force was continuously recorded throughout the test, and the load at which observable crack initiation and subsequent unstable growth occurred was later extracted from the force–time and force–displacement records.

Synchrotron XCT imaging used a monochromatic X-ray beam of 53 keV, with a detector configuration yielding an isotropic voxel size of 3.2 μm. For each loading stage, approximately 1800–2150 projections were acquired over 180° rotation, with an exposure time of 0.05 s per projection, giving a total scan duration of ∼90 s per volume. Projection images were acquired at 2560 × 2568 pixels (32-bit depth).

### 2.3 Image reconstruction and preprocessing

Projection images were reconstructed into 3D volumes using the SAVU pipeline (Kazantsev et al., 2022), producing a consistent field of view that encompassed the notch region and anticipated crack path. Reconstructed 32-bit tomograms were exported as 3D image stacks. For each specimen and loading stage, the greyscale histogram was examined and an intensity window was chosen to maximise contrast within the trabecular region while avoiding saturation of over- and under-exposed voxels. The windowed data were then linearly rescaled and converted to 8-bit to improve the effective greyscale utilisation for subsequent DVC analysis.

To reduce high-frequency noise without blurring trabecular boundaries, a 3D median filter (radius 1 voxel) was applied in ImageJ. Filtered volumes were rigidly aligned so that the long axis of the semi-core and the presumed crack plane were approximately parallel to the image axes. A central region of interest (ROI) containing the notch and anticipated crack path was then cropped to remove edge artefacts and regions outside the gauge length; this ROI comprised approximately 500 central slices along the longitudinal direction, ensuring that the full crack field was contained at all loading stages. Non-specimen structures such as parts of the loading rig or plastic tube were excluded by manual cropping and intensity-based masking. A global greyscale threshold of 125 was applied to remove non-bone voxels, retaining only the trabecular network within the ROI for further analysis (i.e. the trough between greyscale peaks for bone and non-bone in the frequency distribution plot).

For crack visualisation and automated crack detection, the cropped volumes were orthogonally re-sliced to generate two standardised views (Figure 1b–e): a top view, showing the notch mouth and crack plane in plane view; and a right view, showing crack depth and profile through the specimen thickness. Image stacks at each loading stage were exported as 8-bit raw or TIFF volumes for import into the DVC software and subsequent MATLAB post-processing. The full 3D ROI was used as input for DVC (Section 2.4), while the resliced stacks were used for phase congruency–based crack detection (PCCD) and for manual crack length measurements (Section 2.5).

### 2.4 Digital volume correlation

DVC was used to obtain full-field 3D displacement measurements throughout the trabecular network over the course of crack initiation and propagation. Correlations were performed in LaVision StrainMaster DaVis 8.4.0 (LaVision GmbH, Göttingen, Germany), using the dedicated porous-material DVC module optimised for low-density, heterogeneous microstructures (Peña Fernández et al., 2018; Ma, 2018; Cinar, 2019).

For each specimen, the reconstructed volume at preload (Stage 0) was taken as the reference configuration. All subsequent loading stages were correlated back to this reference volume, yielding displacement fields that describe cumulative deformation and crack opening relative to the initial state. A multi-pass, coarse-to-fine strategy was adopted: an initial pass with a large subset size and reduced overlap captured the overall deformation field, followed by several refinement passes with progressively smaller sub-volumes and increased overlap to resolve steep displacement gradients near the crack. Final subset edge length and step size were 64 and 16 voxels respectively (corresponding to approximately 0.205 mm and 0.051 mm), giving an effective spatial resolution of ∼0.05 mm for local displacement and strain measurements. Prior to correlation, a binary ROI mask was generated from the reference volume using the intensity-based masking described in Section 2.3 (global greyscale threshold = 125).

Rigid-body motion between loading stages was corrected following Mostafavi et al. (2015): a large sub-volume encompassing the entire specimen was correlated first to estimate rigid translation and rotation, and the resulting rigid-body transformation was removed from the displacement field before the finer passes. This ensured that the remaining displacements predominantly reflected elastic deformation and crack-induced discontinuities.

Correlation quality was monitored using the normalised cross-correlation coefficient and by visual inspection of displacement and strain maps. Voxels with correlation coefficients below 0.8 or with unphysical displacement outliers were excluded from subsequent analysis using an outlier filter within DaVis and additional masking in MATLAB. To quantify measurement uncertainty, two repeated scans at the same load level were correlated for a representative specimen; the standard deviation of the three displacement components in nominally undeformed regions was ∼0.42 μm, corresponding to less than one-tenth of a voxel and < 1/10 voxel equivalent uncertainty in crack mouth opening displacement (CMOD; defined as the normal displacement separation across the notch mouth; see Section 2.5.1).

### 2.5 Crack parameter extraction

Two complementary approaches were used to quantify crack opening and growth: (i) manual measurements from resliced XCT image stacks, and (ii) automated phase-congruency–based crack detection (PCCD) applied to DVC displacement discontinuities. The combination of 3.2 μm voxel size and the chosen DVC subset spacing imposed a minimum reliably resolvable crack-length increment of approximately 0.05 mm; shorter crack growth segments at the onset of fracture therefore fell below the DVC spatial resolution and were identified manually from the image stacks (Section 2.5.1). PCCD was used to segment the evolving crack surface and quantify crack length at loading stages where the displacement discontinuity was resolvable (Section 2.5.2). Outputs from both approaches were then combined to compute specimen-scale crack-opening descriptors (CMOD/a, (CMOD/a)*, and Δa*; Section 2.5.3).

#### 2.5.1 Manual measurements

Manual measurements were used to quantify crack length and CMOD at each loading stage and were particularly important during the earliest stages of crack development, where incremental crack extension could be below the minimum resolvable length increment (∼0.05 mm) imposed by the DVC subset spacing. For each loading stage, the top-view resliced stack (Section 2.3) was inspected in ImageJ and the crack was identified as a notch-connected discontinuity in the trabecular network emerging from the notch root. Crack length was estimated by counting the number of consecutive slices (Nslices) in which this discontinuity was visible, and converting to a physical length using the voxel size v = 3.2 μm:

a = Nslices · v.

To minimise observer bias, crack visibility was assessed independently by two observers using a fixed scrolling order; where assessments differed, the more conservative (shorter) crack length was adopted. Crack initiation was defined as the first loading increment at which a continuous, notch-connected discontinuity could be followed for at least three consecutive slices in the top-view stack.

CMOD was obtained from the DVC displacement fields as the normal component of displacement separation at the notch mouth. Small rectangular mask windows were placed on the opposing crack faces at the notch mouth, the mean normal displacement within each window was extracted, and CMOD was calculated as the difference between the upper- and lower-face mean displacements. The same procedure was repeated using both a manually defined line of interest and an automatically generated notch-mouth-aligned line to confirm robustness to window placement.

#### 2.5.2 Automated crack detection (PCCD)

Automated crack detection and crack-length quantification were performed using a phase-congruency–based crack detection (PCCD) pipeline applied to the DVC-derived displacement fields, adapted from prior work in quasi-brittle materials and trabecular bone (Cinar et al., 2017; Cinar, 2019). For each loading stage, the DVC displacement field was exported from DaVis (LaVision StrainMaster v8.4.0) as a volumetric vector field (.dat) and imported into MATLAB. The right-view resliced displacement stacks (Section 2.3) were used to standardise crack orientation and facilitate segmentation of displacement discontinuities.

The normal-displacement component was prefiltered using a bilateral filter to suppress high-frequency noise while preserving sharp displacement jumps across crack faces (spatial kernel size = 4; displacement-range parameter = 1.2). A Sobel edge detector was applied to highlight candidate discontinuity edges, and a Hough transform was used to identify the dominant crack trajectory and define a set of seed points along the anticipated crack path. Around each seed, local patches were analysed using a phase congruency transform to detect features based on alignment of local Fourier components, improving robustness to heterogeneous background typical of trabecular architectures. Voxels exceeding the phase-congruency response threshold were classified as candidate crack voxels and used to initialise an active-contour (snake) segmentation that evolved to fit the crack surface while enforcing connectivity and smoothness constraints, allowing tortuous and locally branching crack geometries to be captured.

For each stage, the segmented crack surface was reconstructed in 3D and crack length (a) was measured along predefined cross-sections perpendicular to the notch line. Crack length was evaluated at five equally spaced positions in the longitudinal direction (Y′) across the notch region, and the mean of these five values was taken as the representative crack length for that stage; the standard deviation across positions provided an estimate of crack-path variability and segmentation uncertainty. Segmentation precision was assessed by applying PCCD to repeated scans acquired at the same load level and by comparison with manual crack-length measurements; the resulting uncertainty in crack-front position was approximately ±16 voxels (≈ ±0.051 mm).

#### 2.5.3 Derived parameters

Using CMOD (Section 2.5.1) and crack length a (Sections 2.5.1–2.5.2), crack-opening descriptors were computed at each loading stage. To reduce sensitivity to specimen size, notch depth, and crack-front morphology within the non-standard semi-core geometry, CMOD was normalised by crack length to give the geometry-normalised crack-opening ratio CMOD/a. The critical plateau value (CMOD/a)* was defined as the CMOD/a value at the onset of unstable crack growth (as identified from the mechanical record and corresponding XCT stage). Total crack extension was quantified as Δa*, defined as the increase in crack length from the crack initiation stage (first notch-connected crack meeting the initiation criterion in Section 2.5.1) to the final recorded stage.

### 2.6 Statistical analysis

All statistical analyses were performed at the specimen level, treating each semi-core as an independent observation. For each variable, data were summarised as mean ± standard deviation (SD) for approximately normally distributed variables and as median ± interquartile range (IQR) for skewed variables (from visual inspection of histograms). Hip-Fx and Control specimens were compared using two-tailed Mann–Whitney U tests for crack descriptors including the geometry-normalised crack opening ratio CMOD/a and its critical plateau value (CMOD/a)*, and normalised crack extension Δa*. The association between manual and automated (PCCD-based) crack length measurements was quantified using Pearson’s correlation coefficient (r), computed across all specimens and loading stages where both measurements exceeded the DVC spatial-resolution threshold (∼0.05 mm). All hypothesis tests used a significance level of α = 0.05. Statistical analyses were conducted using GraphPad Prism (GraphPad Software, San Diego, CA, USA).

## 3. Results

### 3.1 Donor characteristics

Donor characteristics are summarised in Table 1. Hip-fracture (Hip-Fx) donors were on average 7.6 years older than Controls (81.4 ± 5.6 vs 73.8 ± 10.7 years; p = 0.208) and had significantly lower BV/TV (∼20%; 0.26 ± 0.03 vs 0.31 ± 0.03; p = 0.019). Sex distribution differed between groups (Hip-Fx 4F/1M; Control 2F/3M).

### 3.2 Crack opening behaviour

CMOD increased nonlinearly with applied load in all specimens. When normalised by crack length (CMOD/a), Hip-Fx specimens showed lower values than Controls across the loading range (Figure 4a). In each specimen, CMOD/a approached an approximately constant plateau, defining the critical crack opening ratio (CMOD/a)*. Hip-Fx specimens reached this plateau at lower applied forces and had significantly lower critical crack opening ratios than Controls: median (CMOD/a)* was 0.31 vs 0.47 (p = 0.008; Figure 4b).

**Figure 4.**
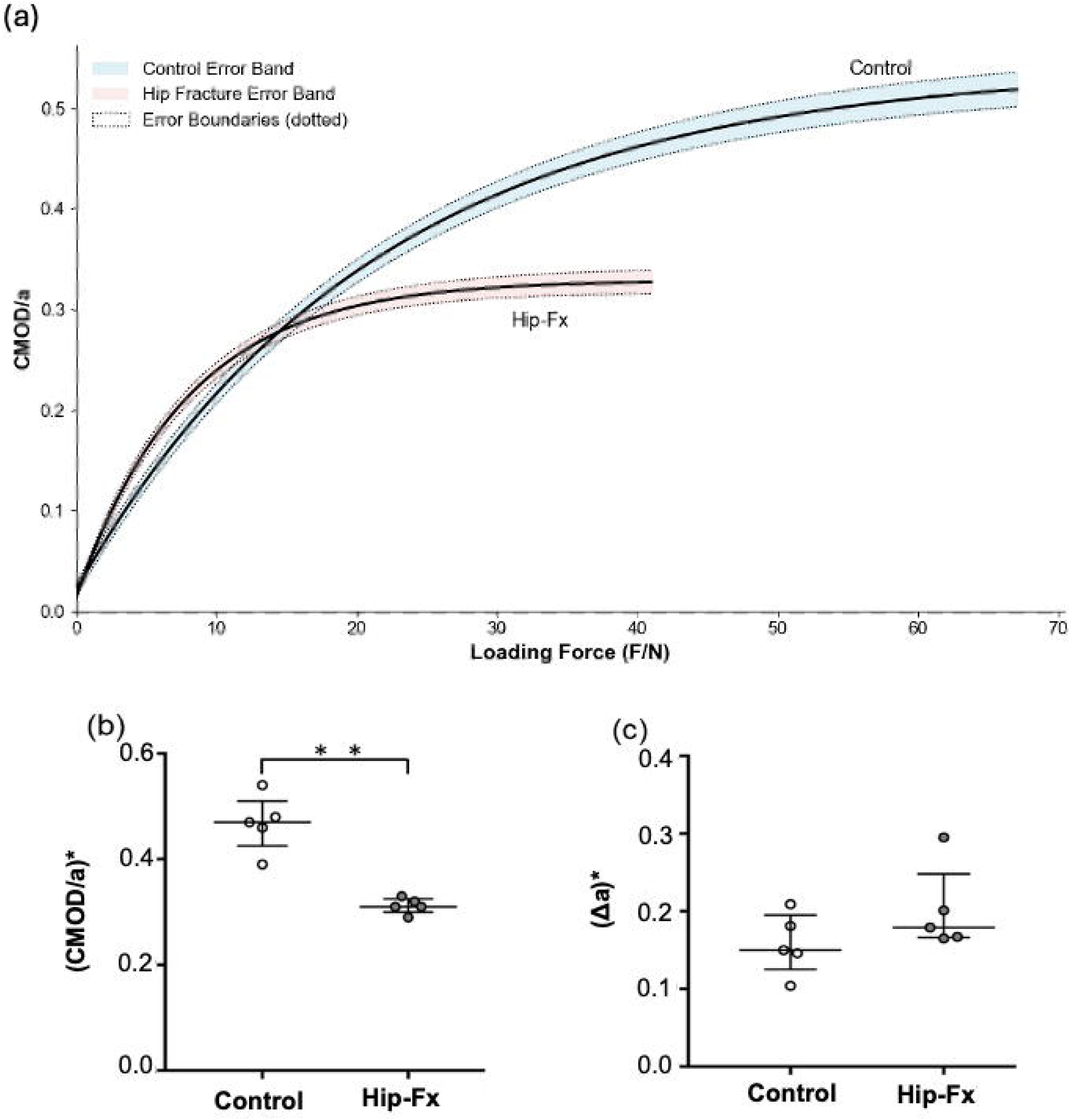
Comparative crack-opening behaviour descriptors for hip-fracture and control specimens. (a) CMOD/*a* versus applied force for Control (*n* = 5) and Hip-Fx (*n* = 5) groups; solid lines indicate mean values and dashed lines denote 95% confidence intervals. (b) Critical plateau values (CMOD/*a*)*, significantly lower in Hip-Fx specimens (p < 0.010). (c) Total crack extension (Δ*a*\*) showing no significant difference between groups (ns). Data presented as median ± interquartile range; Mann-Whitney U test used for comparisons.

### 3.3 Total crack extension

Normalised total crack extension, Δa* (from crack initiation to sudden failure), did not differ significantly between groups (Figure 4c). Median Δa* was 0.179 for Hip-Fx specimens and 0.150 for Controls (p > 0.05). Thus, both groups showed similar overall crack extension before catastrophic failure.

### 3.4 Agreement between manual and automated measurements

Automated crack length measurements from PCCD showed excellent agreement with manual measurements (r² = 0.98; Figure 5). Residual differences remained below the spatial-resolution limit of the DVC analysis (∼0.05 mm) once cracks exceeded the displacement-noise threshold, indicating that PCCD-derived measurements are suitable for calculating CMOD/a, (CMOD/a)* and Δa*.

**Figure 5.**
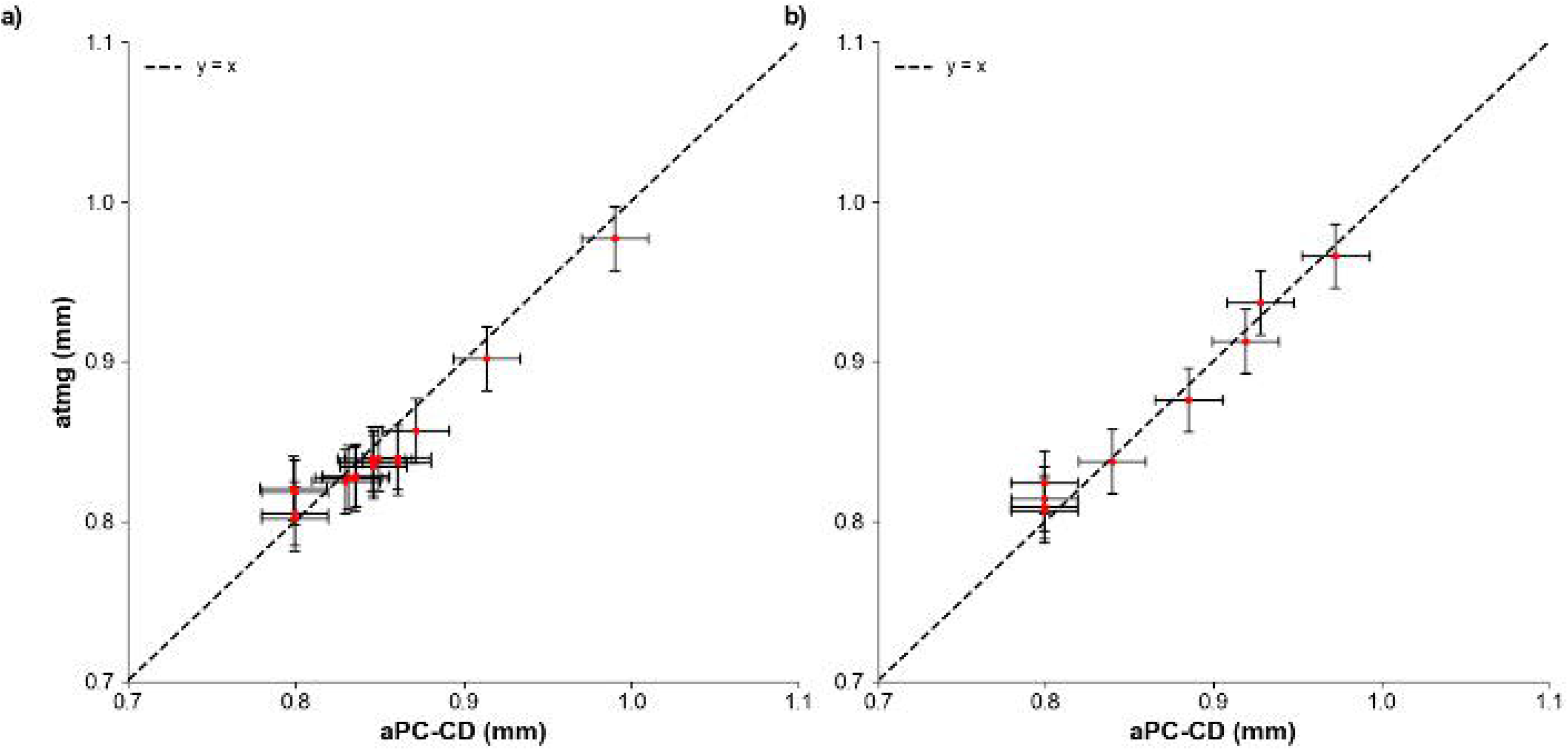
Validation of automated PCCD crack length measurements against manual inspection. (a) Comparison of automated and manual crack length measurements for Control specimens. (b) Corresponding comparison for Hip-Fx specimens. A strong correlation (r² = 0.98) demonstrates high measurement consistency between methods. Vertical error bars indicate standard deviation from five repeated manual measurements; horizontal error bars reflect PCCD precision (±16 voxels ≈ ±0.051 mm). Clustering around ∼0.8 mm corresponds to early increments where crack lengths approached the DVC resolution limit.

## 4. Discussion

This study used synchrotron X-ray computed tomography combined with digital volume correlation (XCT–DVC) to quantify crack-opening behaviour in human femoral-head trabecular bone under three-point bending, using displacement-based, geometry-normalised descriptors in a setting where classical fracture-mechanics parameters cannot be defined reliably. We measured crack mouth opening displacement (CMOD), crack length (a), and their ratio CMOD/a, and combined these with automated phase-congruency–based crack segmentation to track crack evolution through loading. The principal finding was that Hip-Fx specimens exhibited lower critical CMOD/a values and reached instability at lower applied loads than Controls, despite similar total crack extension (Figure 4a–c). Taken together, these findings indicate a more brittle comparative response in Hip-Fx trabecular bone under this test configuration. We therefore present CMOD/a not as a material property or formal toughness parameter, but as a practical comparative descriptor that can be extracted reproducibly from anatomically constrained trabecular specimens.

### 4.1 Comparative brittle response in hip-fracture bone

Hip-Fx specimens exhibited lower critical (CMOD/a)* values and reached instability at lower applied loads than Controls (Figure 4a–b), consistent with a more brittle comparative response and reduced capacity to accommodate deformation before failure. Control specimens showed a more gradual evolution of crack opening, consistent with greater structural robustness. These differences need to be interpreted in the context of the displacement-based descriptors used in this study. Because the semi-core geometry and discontinuous trabecular lattice do not satisfy the requirements for standard fracture-mechanics testing, classical parameters such as K or J could not be evaluated reliably. Instead, crack opening was quantified directly from DVC-derived displacement discontinuities at the notch mouth (CMOD), and normalised by the measured crack length a to form the ratio CMOD/a. This geometry-normalised descriptor reduces sensitivity to specimen size, notch depth and crack-front morphology within the constraints of the semi-core configuration and is used here as a comparative indicator of brittle behaviour rather than an absolute toughness parameter. Accordingly, CMOD/a is interpreted here as a comparative descriptor of crack-opening behaviour, not as an intrinsic material property or a substitute for formal fracture-toughness parameters. Within this approach, the lower critical CMOD/a* in Hip-Fx bone is consistent with reduced tolerance for crack opening relative to crack extension. In other words, Hip-Fx specimens reached instability at lower normalised crack-opening values than Controls under the same experimental configuration. We interpret this as evidence of a more brittle comparative response, while recognising that CMOD/a is not a material property and should not be interpreted as an absolute toughness measure.

### 4.2 Distinct failure pathways despite similar crack extension

Despite these clear differences in critical CMOD/a* and load at instability, total crack extension (Δa*) did not differ significantly between groups (Figure 4c). Both Hip-Fx and Control specimens reached comparable end-stage crack lengths, but they did so via different crack-opening trajectories. In Hip-Fx specimens, CMOD/a rose rapidly and approached its critical value at relatively modest crack extensions, consistent with less opportunity for trabecular bending, crack deflection and bridging to redistribute stresses before instability. In Controls, the evolution of CMOD/a with load was more gradual, suggesting more progressive engagement of local damage-tolerance mechanisms and a greater capacity to sustain crack growth before instability. These observations emphasise that overall crack length alone is insufficient to characterise brittle behaviour in trabecular bone. It is the way crack opening develops relative to crack extension—captured here by the geometry-normalised CMOD/a metric and its critical value—that distinguishes a more brittle Hip-Fx response from a more damage-tolerant Control response under this test configuration.

### 4.3 Agreement between manual and automated crack length measurements

Automated crack detection using the PCCD algorithm showed excellent agreement with manual XCT-based measurements (*r²* = 0.98; Figure 5). This strong correlation supports the use of displacement discontinuity analysis for tracking crack evolution in heterogeneous porous materials (Cinar et al., 2017; Cinar, 2019). Increased variability in manual measurements during early increments reflected limitations imposed by voxel size and DVC subset overlap (Peña Fernández et al., 2018; Ma, 2018). Once displacement discontinuities exceeded these thresholds, PCCD segmentation yielded highly stable and reproducible crack length estimates.

### 4.4 Methodological considerations and limitations

The findings of this study should be interpreted in light of several methodological considerations. This work was primarily designed as a demonstration of the feasibility of applying XCT–DVC to trabecular bone, and the modest sample size (n = 5 per group) therefore limits statistical power; nonetheless, the internal consistency of CMOD-derived descriptors, together with the strong agreement between manual and automated measurements, supports the robustness of the observed trends. Anatomical constraints of the femoral head necessitated a short-span three-point bending configuration, which introduces mixed-mode loading and precludes calculation of classical fracture toughness parameters (Ma, 2018). The experimental configuration was not intended to reproduce physiological loading scenarios; rather, it was chosen to promote controlled, quasi-static crack growth along a well-defined path that is amenable to XCT–DVC measurement. The inability of DVC to capture the very earliest stages of crack initiation is consistent with established detection thresholds determined by voxel size and subset spacing (Ma et al., 2017; Yan et al., 2020). Although XCT entails radiation exposure, uniform imaging conditions across specimens minimise the risk of systematic group bias, and the observed group differences in BV/TV (characteristic of Hip-Fx donors) may contribute to brittle behaviour and warrant further investigation in microstructure-matched cohorts (Table 1). Overall, the present results should be viewed as characterising material- and microarchitecture-dependent brittle response under a specific, well-controlled bending configuration, rather than providing an exhaustive description of trabecular failure across all possible loading modes.

### 4.5 Implications and future research directions

Collectively, these results indicate that XCT–DVC provides a useful experimental approach for studying brittle failure processes in trabecular bone. By quantifying crack opening and propagation using displacement-based metrics rather than relying solely on geometric crack extension, this approach provides a practical way to study fracture behaviour in architecturally discontinuous tissues where classical fracture-mechanics parameters cannot be applied reliably, while complementing existing structural and computational methodologies. In this demonstrator application, displacement-based crack-opening measures resolved differences in brittle behaviour between Hip-Fx and Control donor groups under stringent geometric and loading constraints, suggesting that XCT–DVC can detect group-level variations in comparative brittle response where traditional toughness parameters are not definable. Looking forward, integrating XCT–DVC with higher-resolution imaging, microarchitectural quantification, computational modelling, or machine learning–based microstructure analysis offers a route to multiscale studies aimed at linking trabecular microarchitecture to localised deformation patterns, mechanical resilience, and crack propagation pathways in porous bone.

## 5. Conclusion

Synchrotron XCT combined with DVC enabled direct measurement of crack opening and crack growth in human trabecular bone during stepwise fracture testing. In this anatomically constrained, non-standard configuration, CMOD, crack length, and the ratio CMOD/a provided practical comparative descriptors of brittle response that distinguished Hip-Fx from Control specimens. Hip-Fx specimens exhibited lower critical CMOD/a* values and reached instability at lower applied loads, consistent with a more brittle response under this test configuration, whereas total crack extension did not differ between groups. These findings support the value of displacement-based crack-opening measurements for comparing trabecular fracture behaviour when classical fracture-mechanics parameters are not applicable. The strong agreement between manual and automated crack-length measurements further supports the reproducibility of the measurement pipeline. Overall, XCT–DVC offers a useful experimental approach for comparative studies of crack-opening behaviour in small, architecturally complex bone specimens.

## 6. Acknowledgments

We thank the direct care teams at St Mary’s Hospital for consenting patients and collecting tissue samples. The authors would also like to thank the patients who consented to donate tissue for research and the Imperial College Healthcare National Health Service (NHS) staff and Imperial Tissue Bank staff who helped with the collection of the samples. We thank Vesalius Clinical Training unit and the donors who consented to donate cadaveric tissue for research. We would also like to thank Dr Brian Connolly (University of Manchester) for loaning the in-situ loading rig, provided through EPSRC Grant EP/H025286/1: Long Term, In Situ Material Degradation Studies Utilising High Resolution Laboratory X-ray. We thank Diamond Light Source for access to beamline I12 (JEEP) under proposal EE11204-1, which provided the synchrotron XCT facilities used in this study.

